# Lifestyle impacts the oral microbiome of Classical and Post-classical societies in Italy

**DOI:** 10.64898/2026.01.23.700545

**Authors:** Martina Farese, Markella Moraitou, Chenyu Jin, Adrian Forsythe, Ileana Micarelli, Tom van der Valk, Giorgio Manzi, Laura Parducci, Mary Anne Tafuri, Katerina Guschanski

## Abstract

**Objectives:** The fall of the Roman Empire (476 CE) profoundly affected the lives of its peoples due to the political, administrative, and territorial changes that occurred. The majority of written records of the time focus on the social élite, leaving larger parts of the population understudied. Here, we employ a bioarchaeological approach to understand how differences in lifestyle may be reflected in the oral microbiome of people from different social classes living before and after the fall.

**Material and Methods:** We analysed shotgun sequencing data from dental calculus, the preserved oral microbiome, of 67 individuals belonging to different social classes from two Classical cemeteries dated to I-III century CE (Lucus Feroniae and Isola Sacra) and one post-Classical cemetery dated to IV-VIII century CE (Selvicciola), all located in proximity to the city of Rome, Italy.

**Results:** We detect significant differences in the oral microbiome taxonomic and functional composition across time periods and social classes, with the rural town of Lucus Feroniae standing out compared to its two counterparts. Reliable identification of dietary items was not possible.

**Discussion:** The distinct oral microbiome of Lucus Feroniae could reflect differences in general health and subsistence practices. The rural position of this community may have mitigated the cyclical food crises that, instead, affected the contemporary Isola Sacra and the later community of Selvicciola, thereby buffering against the nutritional stress observed in these two locations.

## INTRODUCTION

Lifestyles in the Roman Empire and Late Antiquity have been extensively researched in the last two decades, with studies aiming to reconstruct social organisations, dietary practices, health conditions and economic activities (Barchiesi & Scheidel, 2010; Hingley, 2005; Purcell, 2003; Twiss, 2019). The surviving written records also offer means to investigate the many facets that constituted Roman and post-Classical societies (approximately 753 BCE - 1492 CE). However, as written sources are usually dominated by the most influential societal strata (the élite), studies based on these records are often biased towards the lifestyles of the upper class who lived in central cities, overlooking the daily lives of the majority of the population. We, therefore, lack a comprehensive understanding of the everyday lives of people from less prominent social classes, who mainly lived outside major cities. This societal stratum is dominated by farmers, war veterans, merchants, freedmen and slaves, who played essential roles in the economy of the times but are underrepresented in historical texts and iconography (Allison, 1999; Dench, 2010; Hingley, 2005; Tacoma, 2016). Direct analyses of skeletal material offer insights into the changes brought about by periods of major social, cultural and political upheaval, as in the case of the fall of the Roman Empire (476 CE). The study of human remains recovered from archaeological sites around the city of Rome can help address the gap left by written records, including uncovering changes in health, diet and living conditions in both rural and urban areas, therefore providing a more complete picture of the past.

Different techniques, such as morphological, palaeopathological, isotopic and molecular analyses, have been applied to investigate the lifestyles, diets and pathologies of ancient peoples, including those from lower societal classes (Emery, Duggan, et al., 2018; Emery, Stark, et al., 2018; Manzi et al., 1991; Salvadei et al., 2001; Tafuri et al., 2018). Recently, the study of dental calculus, the mineralised form of the oral microbiome, has provided unique insights into ancient societies (Bravo-Lopez et al., 2020; Eisenhofer et al., 2020; Gancz et al., 2023; Ottoni et al., 2021; Quagliariello et al., 2022; Velsko et al., 2022, 2023, 2024; Warinner et al., 2014). Dental calculus is formed by the calcification of dental plaque (Y. Jin & Yip, 2002). The mineralised deposits fossilise on the living host, reducing post-mortem contamination from exogenous microorganisms and allowing the preservation of endogenous DNA in a virtually unchanged form for millennia (Mann et al., 2018; Warinner et al., 2015). Dental calculus preserves the remains of oral and respiratory microbial communities, dietary elements and host-derived molecules (Ozga et al., 2016; Velsko et al., 2022; Warinner et al., 2014).

The dental calculus microbiome appears to have been rather stable over time (within the last 8,000 years), even during major cultural transitions, including the introduction of agriculture and the Industrial Revolution (Gancz et al., 2023; Ottoni et al., 2021; Velsko et al., 2023). Several studies have attempted to use the dental calculus microbiome to shed light on how communities and their lifestyles have developed. However, Ottoni et al. (2021) found that the oral microbiomes of humans living in the Palaeolithic and Mesolithic periods remained stable, with only minor changes detected since the introduction of agriculture. Similarly, a comparative study spanning from the Neolithic to the Industrial Revolution found no significant changes in the dental calculus microbiome composition (Gancz et al., 2023). A strong stability was also observed when comparing foraging and highly industrialised populations, with dietary differences playing only a minor role in the composition of dental plaque in human populations with different subsistence strategies (Velsko et al., 2023). However, despite a lack of temporal changes, a distinct oral microbial community composition was recovered from dental calculus in Oceania, setting it apart from Europe, Asia and Africa (Velsko et al., 2024). Supporting this, the dental calculus microbiome composition and function of closely related gorilla (sub)species were shown to vary with host species ecology, most likely reflecting different diets (Moraitou et al., 2022), despite their geographic proximity and recent divergence of only 12,000 years ago (Van Der Valk et al., 2024). This suggests that subtle but persistent differences in lifestyle and diet might be reflected in the dental calculus microbiome in humans as well. If so, this will open doors to studying ancient human communities that lack other sources of information about their lifestyles.

So far, only a few studies have examined human communities living in geographic proximity but differing in their lifestyles and diet (Cameron et al., 2015; Sarkar et al., 2017; Velsko et al., 2023). However, these studies focused primarily on modern populations, with a notable exception of the recent investigation of archaeological samples from Oceania (Velsko et al., 2024). Here, we aim to advance our understanding of past societies, particularly of communities less represented in classical literature. To do so, we conduct comparative analyses of the dental calculus oral microbiome from three archaeological sites from the vicinity of the city of Rome, Italy (Figure 1), housed in the Museum of Anthropology “Giuseppe Sergi”, at Sapienza University of Rome. These include two contemporaneous communities that lived during the Roman Imperial period but differed in their lifestyles (Isola Sacra and Lucus Feroniae), and a community dating to the Late Antique and Longobard period (Selvicciola). By characterising oral microbiomes, we aim to investigate how differences in socio-economic status during the Roman Empire and the cultural transition that followed its fall impacted local human societies.

**Figure 1.**
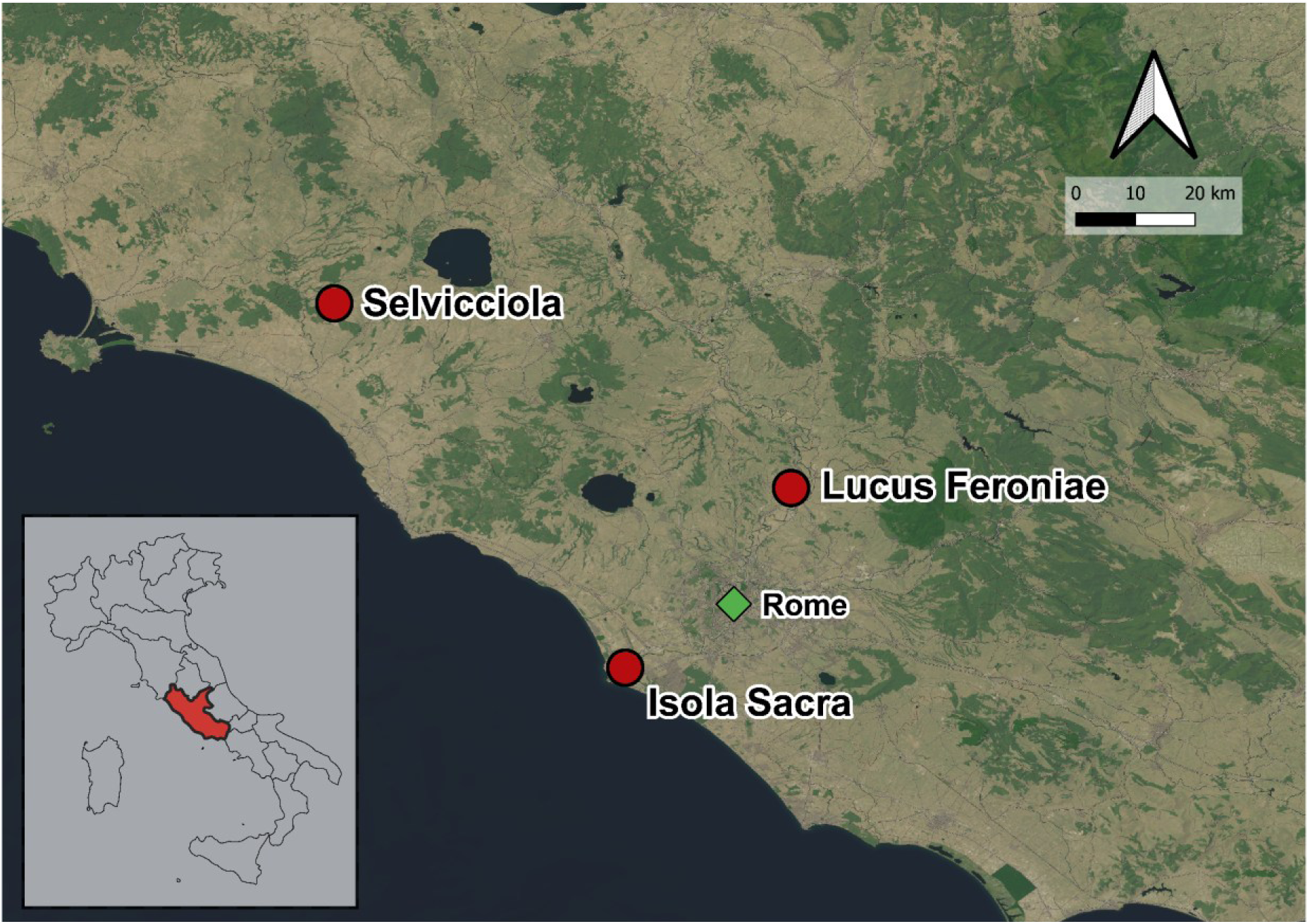
Maps with the location of the archaeological sites presented in this study (red circles), with the city of Rome displayed as a green diamond (map created with QGIS 3.36.3).

## MATERIALS AND METHODS

### Archaeological sites

#### Isola Sacra

The cemetery of Isola Sacra is linked to the ancient city of Portus Romae, about 20km west of Rome (Figure 1). It dates back to the Roman Imperial age and, as no radiocarbon dates are available, the individuals are generally attributed to the I-III century CE, based on the archaeological material recovered. The cemetery is located along the road connecting the city of Portus to the coast and the town of Ostia, the *Via Flavia*, on the so-called “sacred island” (Baldassarre, 2002; Prowse et al., 2004; Verduchi, 1992). Portus was built around the port established by Emperor Claudius and expanded by Emperor Trajan between 42 and 64 CE. The importance of Portus grew during the expansion of Rome, being the main harbour of the capital of the Empire. The inhabitants of Portus included merchants, administrators and traders; however, no local aristocracy is mentioned in the written records. Dietary analysis by means of carbon and nitrogen stable isotopes, performed on a subgroup of 105 individuals buried in the cemetery of Isola Sacra, suggests a mixed diet composed of marine and terrestrial resources (Prowse et al., 2004, 2005).

#### Lucus Feroniae

The cemetery is connected to the Roman town of Lucus Feroniae, 30km northeast of Rome (Figure 1). No radiocarbon dates are available, but, as for Isola Sacra, the cemetery was dated to the I-III century CE on the basis of the archaeological material. Lucus Feroniae was a rural centre with an economy based on wheat cultivation (Gazzetti, 1986). In 31 BCE, after the defeat of Marcus Antonius in Actium, Augustus assigned several areas of the town to his troops, thereby making the soldiers part of the population, alongside the locals (Stanco, 1997). The general lack of grave goods inside the tombs and the presence of developmental and occupational stress markers on the skeletal remains (Manzi et al., 1989; Sperduti, 1997) of several individuals suggest that the cemetery holds manual workers and people of humble origins, war veterans, *liberti* (freedmen and freedwomen) and slaves. The stable isotope data suggest that, despite being a rural town, people living in Lucus Feroniae had access to a variety of food resources, including marine products, as a result of both agricultural activities carried out at the site and the exploitation of the local environment (Tafuri et al., 2018).

#### Selvicciola

The cemetery of Selvicciola, located near the town of Ischia di Castro (Viterbo), 90km north of Rome (Figure 1), has been dated to between the end of the IV and the beginning of the VIII century CE (Gazzetti, 1995, 1997; Incitti, 1997; Micarelli, 2020; Patera, 2008). After the fall of the Western Roman Empire in 476 CE, the area was dominated by the Longobards, with archaeological findings also testifying to the presence of a wealthy community since the VI-VII centuries CE. Selvicciola, a Longobard site dated to the mid-VII century CE, was built over an older Roman settlement (Patera, 2009). The anthropological investigations performed on the human remains from Selvicciola suggest a lower quality of life compared to Lucus Feroniae and Isola Sacra, with a higher incidence of dental and skeletal pathologies (Manzi et al., 1999; Salvadei et al., 2001). The low variability in nitrogen stable isotope data suggests that the diet of individuals buried in Selvicciola was based on a limited selection of resources, with a stronger reliance on grains than in Lucus Feroniae (Tafuri et al., 2018).

### Sample collection

We collected a total of 71 dental calculus samples from 67 individuals housed in the Museum of Anthropology “Giuseppe Sergi”, part of the Sapienza University of Rome Museum Network (*Polo Museale*), from both healthy and carious teeth. Thirty of the sampled individuals were buried in the cemetery of Isola Sacra, 27 in Lucus Feroniae and 10 in Selvicciola (Table S1). Among the collected samples, 58 were from individuals without signs of dental caries (although some showed other oral pathologies, such as antemortem tooth loss, linear enamel hypoplasia, dental fractures or heavy tooth wear), six were from within the caries lesions, and seven were from healthy teeth in individuals that showed caries elsewhere in the oral cavity. We also collected eight soil samples (seven from Lucus Feroniae and one from Selvicciola) to act as environmental controls. As no soil was purposefully sampled during the excavation, these samples were obtained from soil particles that adhered to the bones of the study individuals. We prioritised skeletal positions as far away from the cranium and the dental calculus as possible (Table S1). No soil samples were available from Isola Sacra. Sampling was carried out following procedures specific for ancient DNA sample collections (Sabin & A Fellows Yates, 2020; Velsko et al., 2017; Warinner et al., 2019). All samples were then transferred to Uppsala University, Sweden, for processing.

### Sample preparation and shotgun sequencing

All samples were processed in the dedicated ancient DNA laboratory. The calculus samples (weighing between <5 and 20 mg) were ground with sterile micro-pestles. Surface decontamination was performed as in Brealey et al. (2021), exposing the samples to UV light (245 nm) for ten minutes, followed by a wash in 500 μl of 0.5 M EDTA pH 8 for 30 s. Both dental calculus and soil samples were extracted with a silica-based method (Dabney et al., 2013) adapted for dental calculus, as previously described in Brealey et al. (2020). Briefly, after the wash step and centrifugation, the residual pellets were treated with 1 mL of extraction buffer (0.45 M EDTA pH 8, 0.25 mg/mL Proteinase K) and left to digest in a rotating incubator at 37°C overnight. After centrifugation, the supernatant was mixed with 400 μL of 3 M sodium acetate and 10 mL of binding buffer (5M GuHCl, 40% isopropanol, 0.05% Tween-20) and transferred into silica columns. Washing was performed with 450 μL of PE buffer and DNA eluted with 45 μL of EBT buffer (10 mM Tris-HCl pH 8.5, 0.05% Tween-20). Two extraction blanks, positioned as the first and last tubes, were included in every extraction batch of up to 16 samples to control for laboratory and reagent contamination.

Following DNA extraction, double-stranded genomic libraries (Meyer & Kircher, 2010) were constructed using 20 μL of the DNA. We used the double-barcoding double-indexing strategy (Van Der Valk et al., 2020) to identify and later remove instances of index hopping. One library blank was included in every batch of up to 16 samples. The appropriate number of indexing cycles was estimated using a quantitative PCR assay and ranged from 12 to 20 (Table S1). Qiagen MinElute columns were then used to clean up the libraries, and size selection was performed with Agencourt AMPure XP PCR Purification beads (Beckman Coulter, IN, USA) to remove adapter dimers and fragments above 500 bp. The libraries were pooled (using 1.5 µL of each negative control and 3 µL of each dental calculus and soil sample) and submitted to SciLifeLab (Uppsala, Sweden) for shotgun sequencing on two lanes of an S4 flow cell using the NovaSeq 6000 system and v1.5 sequencing chemistry (150bp PE).

### Data Processing

After demultiplexing, we processed the sequences using the nf-core/eager pipeline (Fellows Yates, Lamnidis, et al., 2021) (v. 2.4.4, available at https://nf-co.re/eager/2.4.4), which performs read merging and adapter removal, filtering, mapping to the human genome and taxonomic classification of unmapped reads using Kraken2. We kept the default parameters, except for the minimum read length after adapter clipping and read merging (adjusted to 30 bp) and the minimum base quality for trimming off bases (30) after adapter removal and read merging. Unmerged reads were excluded from downstream analysis. For BAM filtering, the minimum mapping quality for reads (30) and the minimum read length to be kept after mapping (35 bp) differed from the nf-core/eager default parameters. The mapping against the human reference genome (GRCh38) was performed in the nf-core/eager pipeline using BWA-aln (Li & Durbin, 2009) and unmapped reads were converted to FASTQ format and used downstream for the microbial and dietary analyses. Reads mapping to the human genome were then used for molecular sex assignment.

### Taxonomic analysis

Taxonomic classification was carried out using the standard database of Kraken2 (Wood et al., 2019), which includes all the bacteria, archaea and virus genomes available on NCBI. Bracken (Lu et al., 2017) was used to estimate abundances at the species level, and the *taxize* R package (Chamberlain & Szöcs, 2013) was used to retrieve taxonomic information. Microbial species abundances, taxonomic information and sample metadata were analysed jointly using the *phyloseq* R package (McMurdie & Holmes, 2013).

We performed source tracking with the *FEAST* R package (Shenhav et al., 2019) using reference metagenomes from seven source environments: human dental calculus (N=5), human dental plaque (N=4), human caries (N=3), human gut (N=4), human skin (N=5), common laboratory contaminants (N=4), and soil from Northern Italy (N=4) (Table S2). Dental calculus, plaque and caries were jointly considered as the oral environment for source tracking. We retained dental calculus samples with at least 30% oral content according to the FEAST results.

We then performed abundance filtering by converting any relative abundances accounting for less than 0.01% for a given sample to zero. As some of the microbial taxa found in dental calculus are likely contaminants from the burial environment, we used information from co-extracted soil samples to identify them. Specifically, taxa with a higher relative abundance in any single soil sample than in any dental calculus sample were considered to be derived from the soil microbiome and removed from dental calculus samples and blanks as environmental contaminants.

To assess alpha diversity, we produced rarefaction curves (with subsample size ranging from 100 to 10,000 reads) using the ‘plot_alpha_rcurve’ function from the *microbiomeutilities* R package (Shetty & Lahti, 2020). Following this analysis, we excluded samples with too few reads that were far away from the species richness asymptote, as indicated by the rarefaction curves. We then tested for the effect of cemetery provenance on the observed species richness and the Shannon evenness index of healthy teeth using type II (hierarchical) ANOVA (Analysis of Variance). Although only healthy teeth were analysed, some included individuals had caries elsewhere in the oral cavity. Therefore, individual health state and sample read count (after abundance re-estimation with Bracken) were included as explanatory variables to account for potential effects. We used Shapiro-Wilk tests to assess whether the ANOVA residuals meet the normality assumption.

To investigate the composition of the oral microbiome, we conducted all analyses with taxonomic assignments at the species level and applied a centered-log-ratio (CLR) normalisation using the *microbiome* R package (Lahti & Shetty, 2018). We used Aitchison distances (Aitchison, 1982), which are defined as the Euclidean distances between CLR-transformed abundances. We performed permutational analysis of variance (PERMANOVA) only on healthy teeth and assessed the marginal effects of sample read count, cemetery and individual health state using the *vegan* R package (Oksanen et al., 2011). For any significant factor, we performed pairwise PERMANOVAs using the ‘pairwise.adonis’ function from the *pairwiseAdonis* R package (Martinez Arbizu, 2020). We identified differentially abundant taxa while accounting for unequal sequencing depth across metagenomes with the *ANCOMBC2* R package (Lin et al., 2022; Lin & Peddada, 2020), limiting the test to taxa present in at least 10% of samples and with at least 1000 reads. We adjusted the *p* values for multiple testing using the Benjamini-Hochberg method. Principal Coordinate Analysis (PCoA) plots were generated from CLR-transformed data with the *phyloseq* and *ggplot2* R packages (Wickham, 2016).

### Functional analysis

Functional analysis was carried out using HUMANn3 (Beghini et al., 2021). We used the relative abundance of gene families normalised to copies per million, which were aggregated into GO terms to obtain the functional profiles of the metagenomes (Ashburner et al., 2000; The Gene Ontology Consortium et al., 2021). We only retained GO terms reflecting biological processes (“BP”) and normalised abundances using a CLR transformation. We then visualised the functional composition of soil and dental calculus microbiomes using PCoA ordinations implemented in the *phyloseq* package. Dental calculus samples clustering with soil were considered poorly preserved and removed. The retained dental calculus samples were used to test for functional differences among the three populations, using PERMANOVA with Aitchison distances and controlling for sequencing depth (number of reads per sample). For any significant factors, we performed between-level comparisons using the ‘pairwise.adonis’ function from *pairwiseAdonis*, and differential analysis using *ANCOMBC2* (this time using a prevalence cutoff of 20%).

### Sex assignment

Molecular sex identification was performed using sexassign (Gower et al., 2019). Following mapping to the human reference genome as part of eager, we filtered the resulting BAM files (mapping quality ≥ 30, length ≥ 35bp) and removed reads that mapped to unplaced scaffolds (using samtools view to extract each chromosome and samtools merge to combine them) (Li et al., 2009). We generated coverage statistics using samtools idxstats (Li et al., 2009) as input for sexassign, which uses a probabilistic approach based on the ratio of reads mapping to the X chromosome versus the autosomes to identify the sex of the sampled individuals. A likelihood ratio near 0.5 corresponds to males, a ratio near 1.0 corresponds to females, while sex cannot be assigned for ratios between 0.6 and 0.8 (Gower et al., 2019).

### Inferring likely geographic origin of the study individuals

To obtain insights into the genetic affinities of the study individuals, we first constructed a BWA index for the human reference genome hg19. Next, we aligned the sequence reads to this reference using BWA-aln with settings optimised for ancient DNA (-l 16,500 -n 0.01 -o 2) and randomly sampled an allele at each site where reads mapped using ANGSD (-dohaplocall 1 - doCounts 1 - remove_bads 1 -uniqueOnly 1 - minMapQ 20) (Korneliussen et al., 2014). We then used 2,506 human genomes available in the 1000 Genomes project phase 3 (Sudmant et al., 2015; The 1000 Genomes Project Consortium et al., 2015) and identified variable positions in this reference dataset that were also covered by at least 1 read in our data. To identify the genetically closest reference population for each study individual, we counted the number of matching alleles between our samples and each of the 2,506 human reference genomes. The population with the fewest mismatches per variable site was considered to be the closest present-day population to the ancient samples.

### Dietary analysis

To obtain dietary insights from the dental calculus samples, we classified the sequencing reads to plants and animals. First, we constructed genome-wide databases for plants and mammals using Kraken2 (Table S3). The Kraken2 plant database was built by sampling 1257 NCBI genome accessions of different plant species from different genera, all under the family *Viridiplantae*, all Refseq mitochondrial genomes available in April 2023 and all assemblies available in the PhyloNorway database, which contains skimmed genomes of 1,323 species (retrieved 2023.4.13). In total, this database contained 2,638 plant species from 1,087 genera, of which phylonorway and NCBI assemblies together contribute 2,376 species from 960 genera (C. Jin et al., 2025). A second Kraken2 database for mammals was built using 740 genomes of mammalian species belonging to 412 genera. We classified all merged reads against both these databases in Kraken2 with default settings, using the --report-minimizer-data parameters to obtain the number of unique kmers assigned to each species. Additionally, since classical literature and modern studies point to the consumption of marine resources by these populations, we built a bowtie2 index for 27 fish genomes (Table S3) that could have been consumed at the time (Craig et al., 2009; Locker, 2007; Marzano, 2013, 2018; Purcell, 1995). We then competitively mapped all the merged reads against this single bowtie index using bowtie2, with the –very-sensitive setting, and subsequently used samtools depth to count the number of bases covered for each of the fish genomes.

Reliable identification of eukaryotic sequences, particularly of dietary nature, from dental calculus is challenging (Fagernäs et al., 2022; Mann et al., 2023). We, therefore, implemented several pre-processing steps to reduce the detection of false positives. Specifically, we only retained eukaryotic taxa with at least 100 reads (or at least 100 mapped sites for fish) in dental calculus samples, and used soil samples as environmental controls, excluding taxa as likely environmental contaminants if the average relative abundance in soil samples (or negative laboratory controls) was higher than the average relative abundance in dental calculus samples. Because Isola Sacra lacked soil samples and only a single soil sample was available from Selvicciola, we combined the information from all available soil samples for this analysis.

## RESULTS

### Data Processing, Validation and Alpha Diversity

We collected 71 dental calculus and eight soil samples from three cemeteries in Central Italy (Table 1). Our dataset also included 16 negative controls (including extraction and library preparation negatives) that were processed alongside the samples (Table S1).

**Table 1.** Overview of the samples used in this study. See Table S1 for more details.

| Cemetery | Time period | Latitude (N) | Archaeological period | Longitude (E) | N of dental calculus samples | N unique individuals | N of soil samples |
| --- | --- | --- | --- | --- | --- | --- | --- |
| Isola Sacra | I - III century | 41.769 | Classical | 12.264 | 30 | 30 | - |
| Lucus Feroniae | I - III century | 42.130 | Classical | 12.597 | 30 | 27 | 7 |
| Selvicciola | IV - VIII century | 42.501 | Post-Classical | 11.679 | 11 | 10 | 1 |
| **TOTAL** |  |  |  |  | **71** | **67** | **8** |

We obtained a mean of 66,285,548 paired reads per dental calculus sample (ranging from 4,141,846 to 144,842,179), a mean of 43,495,561 paired reads per soil sample (6,448,503 - 99,467,428) and a mean of 171,139 paired reads from the negative controls (654 - 1,160,363). Thus, negative controls produced at least an order of magnitude fewer reads than samples.

We taxonomically classified the samples using the standard Kraken2 database (Wood et al., 2019) and identified a total of 1,953 microbial species (1,902 of these found in dental calculus, 1,506 in soil and 107 in blanks). Source tracking suggests that, on average, 56.6% of dental calculus metagenome composition resembled the human oral microbiome (range 6.8-89.8%), whereas this fraction was 22.9% for soil (range 0.2-39.6%, Figure 2). Dental calculus samples with less than 30% oral proportion were considered to be poorly preserved and, accordingly, nine such samples were excluded from downstream analysis (Table S1).

**Figure 2.**
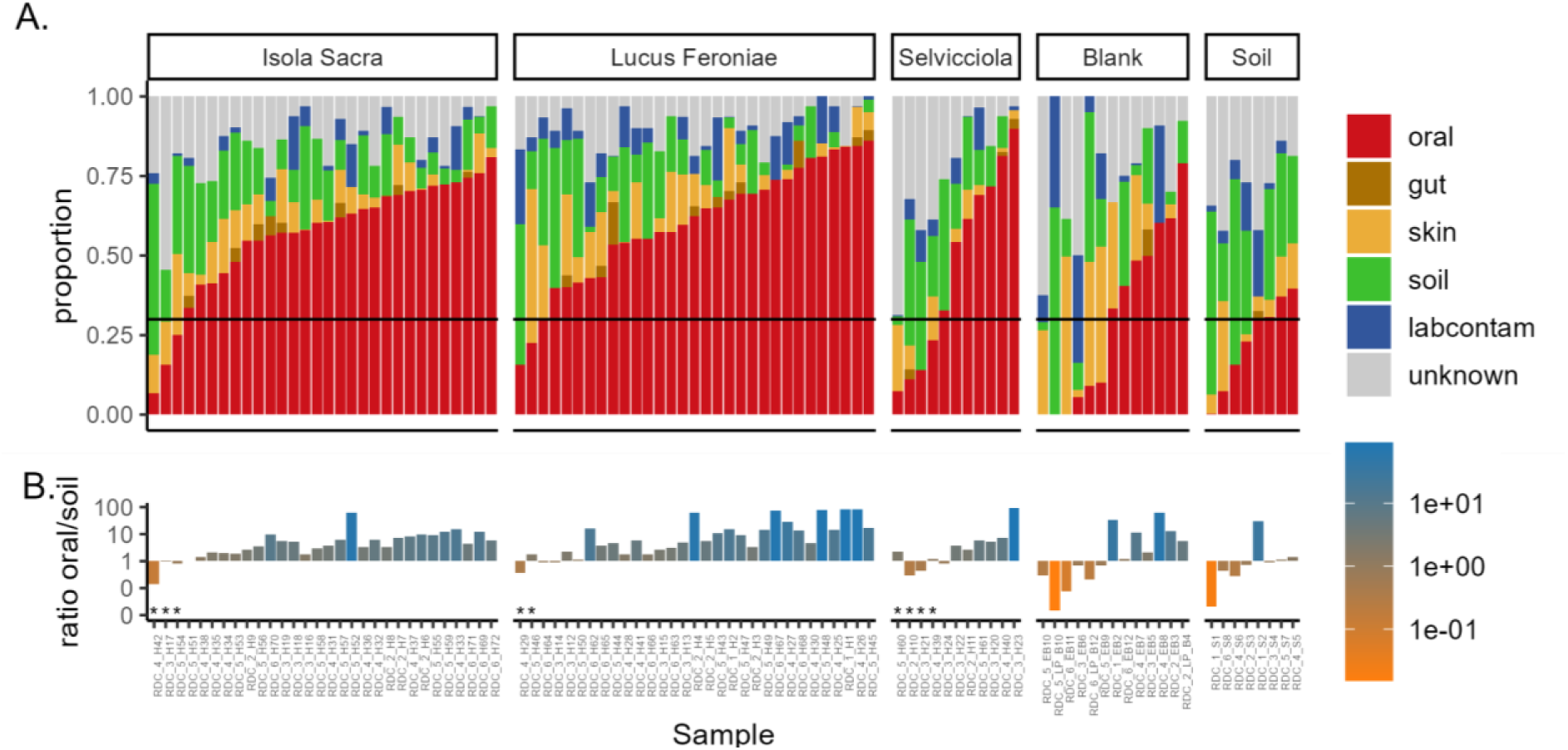
FEAST output for the complete dataset before removing samples with poor preservation and likely contaminant taxa. **A.** Estimated proportion of samples resembling the oral, gut or skin microbiome, common laboratory contaminants and soil. The “oral” category includes human dental calculus, dental plaque and caries sources. The “unknown” category includes taxa that are not part of the microbiome of any of the included sources. Black line at 30% denotes the cut-off for human oral microbiome proportion, below which samples were deemed to be poorly preserved and removed from further analyses. **B.** Ratio of oral to soil proportions for the samples in panel A (note log-transformed y-axis). Pseudo-counts of 0.01 were added to both proportions. Samples with asterisks were excluded due to low preservation (see also Table S1).

By comparing the relative abundances of taxa in dental calculus and in soil, we removed 452 likely environmental contaminants. These included 164 known laboratory contaminants (Salter et al., 2014; Weyrich et al., 2019), six of which are also oral taxa (*Staphylococcus epidermidis*, *S. pettenkoferi, S. pseudoxylosus*, *S. cohnii, Streptococcus pyogenes* and *Actinomyces oris*) (Chen et al., 2010; Fellows Yates, Velsko, et al., 2021). After removing these taxa from dental calculus and blank samples, and performing abundance filtering with a 0.01% relative abundance threshold, we retained 1,362 taxa in the complete dataset (910 of these found in dental calculus, 1,025 in soil and 66 in blanks; Table S4). Following this decontamination procedure, dental calculus, soil and blanks formed separate clusters in a PCoA ordination using Aitchison distances (Supplementary Figure S1).

Rarefaction curves suggest that the number of identified microbial taxa in a dental calculus sample reaches an asymptote after roughly 4,000 taxonomically classified reads (after filtering and decontamination) (Supplementary Figure S2). For this reason, samples with fewer than 4,000 reads were not considered in ANOVA tests assessing microbiome species richness and evenness (Supplementary Figure S3, Table 2, Table S1). The evenness test met the normality of the residuals assumption (Shapiro-Wilk: W = 0.98, *p* value = 0.56), whereas the species richness test had slightly non-normal residuals (W = 0.94, *p* value = 0.016). We did not detect any differences in species richness or evenness by cemetery, although read count affected evenness, even after excluding samples with too few reads (Table 2).

**Table 2.** Species richness and evenness of dental calculus microbiome of healthy teeth do not differ across cemeteries after accounting for read count and health state (i.e., presence of caries elsewhere in the oral cavity). Results of ANOVA tests with Type II sums of squares for observed species richness and Shannon index.

| Response | Observed species richness |  | Shannon index |  |
| --- | --- | --- | --- | --- |
|  | F-statistic | p value | F-statistic | p value |
| Read count | 1.820 | 0.185 | 17.130 | <0.001 |
| Individual health state | 1.040 | 0.314 | 1.640 | 0.200 |
| Cemetery | 1.410 | 0.254 | 0.410 | 0.660 |

### Oral microbiome composition and function are associated with differences in lifestyle

To investigate whether the differences in social status and time period that characterise the three cemeteries affected the oral microbiome, we compared microbial compositions across the cemeteries using PERMANOVA with Aitchison distances (on 58 samples, excluding the four caries samples). We estimated the marginal effects of sequencing depth, individual health state and cemetery provenance, and observed only a small but statistically significant effect of cemetery provenance on the dental calculus bacterial communities (Table 3, Figure 3A). Specifically, we found that the Classical period cemetery of Lucus Feroniae differs significantly from both the contemporaneous cemetery of Isola Sacra and the post-Classical cemetery Selvicciola (Table 3, Figure 3A).

**Figure 3.**
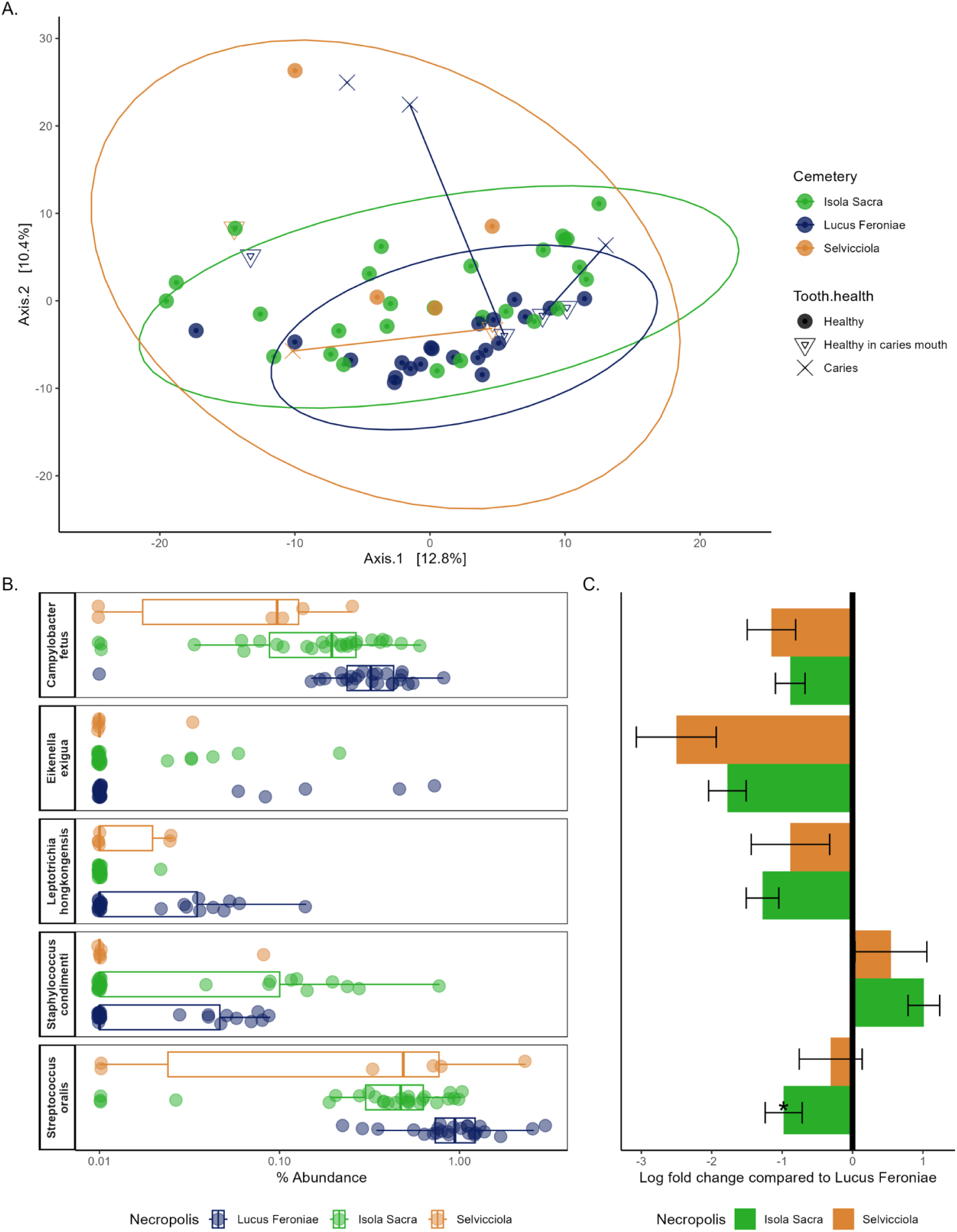
**A.** Compositional differences between 62 dental calculus samples that were retained after filtering and decontamination: 27 from Isola Sacra (in green), 28 from Lucus Feroniae (in blue) and 7 from Selvicciola (in orange). Four caries samples (cross symbols) and six samples of healthy teeth in individuals with caries (triangle symbols) are included. Three pairs of healthy-caries teeth from the same individual are connected by a line. The ellipses show the 95% confidence level using a multivariate t-distribution. **B.** Relative abundances of differentially abundant microbial species between Lucus Feroniae and Isola Sacra, identified using *ANCOMBC2* with Lucus Feroniae used as the reference. Note the log-transformed (with 1% pseudo-count) x-axis. **C.** Log-fold taxon abundance changes compared to Lucus Feroniae for the microbial species in B, as estimated by *ANCOMBC2*. The error bars reflect the standard error of the log-fold change. The asterisk highlights *S. oralis*, which passed the *ANCOMBC2* sensitivity analysis for pseudo-count addition in Isola Sacra.

**Table 3.** Compositional differences across cemeteries (PERMANOVA and pairwise PERMANOVA using Aitchison distances) showing the effect size (R^2^) and *p* value of read count (as a control for sequencing depth), health state (presence of caries in the sampled individual), and cemetery provenance for the 58 dental calculus samples collected from healthy teeth.

| PERMANOVA terms | SumOfSqs | $R^2$ | $p$ value |
| --- | --- | --- | --- |
| Read count | 621.100 | 0.020 | 0.217 |
| Individual health state | 532.100 | 0.017 | 0.383 |
| Cemetery | 1730.800 | 0.056 | 0.005 |
| Residuals | 27831.100 | 0.902 |  |
| Pairwise PERMANOVA | SumOfSqs | $R^2$ | Adjusted $p$ value (Benjamini-Hochberg) |
| Isola Sacra vs Lucus Feroniae | 925.000 | 0.035 | 0.007 |
| Isola Sacra vs Selvicciola | 747.320 | 0.038 | 0.166 |
| Lucus Feroniae vs Selvicciola | 1088.580 | 0.074 | 0.007 |

We were able to compare dental calculus obtained from a healthy and a carious tooth for three individuals. The microbiomes from carious teeth appear more dispersed compared to their healthy counterparts, a pattern also observed in an additional caries sample from Lucus Feroniae without a healthy pair (Figure 3A). Although the sample size is too small to draw general conclusions, caries samples included here do not appear to show a directional shift and do not cluster together, being instead positioned on the margins of the PCoA plot. This could potentially reflect their variable microbiome composition.

Since Lucus Feroniae differed from both Isola Sacra and Selvicciola, we used it as the reference group in the differential abundance analysis with *ANCOMBC2*. Although Lucus Feroniae and Selvicciola showed the most marked difference in microbiome composition (Table 3, Figure 3A), we were unable to identify any differentially abundant microbial species between the two cemeteries. This could be due to the low sample size of Selvicciola (N=7 retained samples). However, we identified five taxa that differ in abundance between the contemporaneous cemeteries of Lucus Feroniae and Isola Sacra. *Campylobacter fetus, Eikenella exigua*, *Leptotrichia hongkongensis* and *Streptococcus oralis* were enriched in Lucus Feroniae, and *Staphylococcus condimenti* was enriched in Isola Sacra (Figure 3B,C). *L. hongkongensis* and *S. oralis* are included in the Human Oral Microbiome database (HOMD) (Chen et al., 2010), whereas the remaining three species are not part of HOMD, but other species belonging to the same genera are. *E. exigua, L. hongkongensis* and *S. condimenti* were found in only 20.7%-36.2% of samples and, when present, showed low average relative abundances ranging from 0.03% (*L. hongkongensis*) to 0.15% (*E. exigua*). *C. fetus* and *S. oralis* were present in 89.7% and 91.4% of samples and had an average abundance of 0.27% and 0.81%, respectively (Table S4). Only *S. oralis* passed the *ANCOMBC2* sensitivity analysis, being consistently identified as differentially abundant regardless of the pseudo-count used by the software.

To understand whether the three populations differed in the function of their oral microbiomes, we compared the compositions of Gene Ontology (GO) terms referring to biological processes. We visualised these functional compositions for both soil and dental calculus microbiomes using PCoA and identified ten dental calculus samples that clustered with soil, indicating poor preservation (Supplementary Figure S4). As GO terms are often encoded by multiple microbial taxa, decontamination based on taxon abundance, as performed for taxonomic analyses, was not possible. Instead, we removed these ten samples from downstream analysis, retaining 52 dental calculus samples (22 from Isola Sacra, 25 from Lucus Feroniae, 5 from Selvicciola) and 1,099 GO terms. After accounting for differences in sequencing depth, PERMANOVA (without caries samples) showed a significant difference in microbial functions across cemeteries (Figure 4A, Table 4, Table S5). As was the case for the taxonomic characterisation, Lucus Feroniae was significantly different from the other two cemeteries (Table 4).

**Figure 4.**
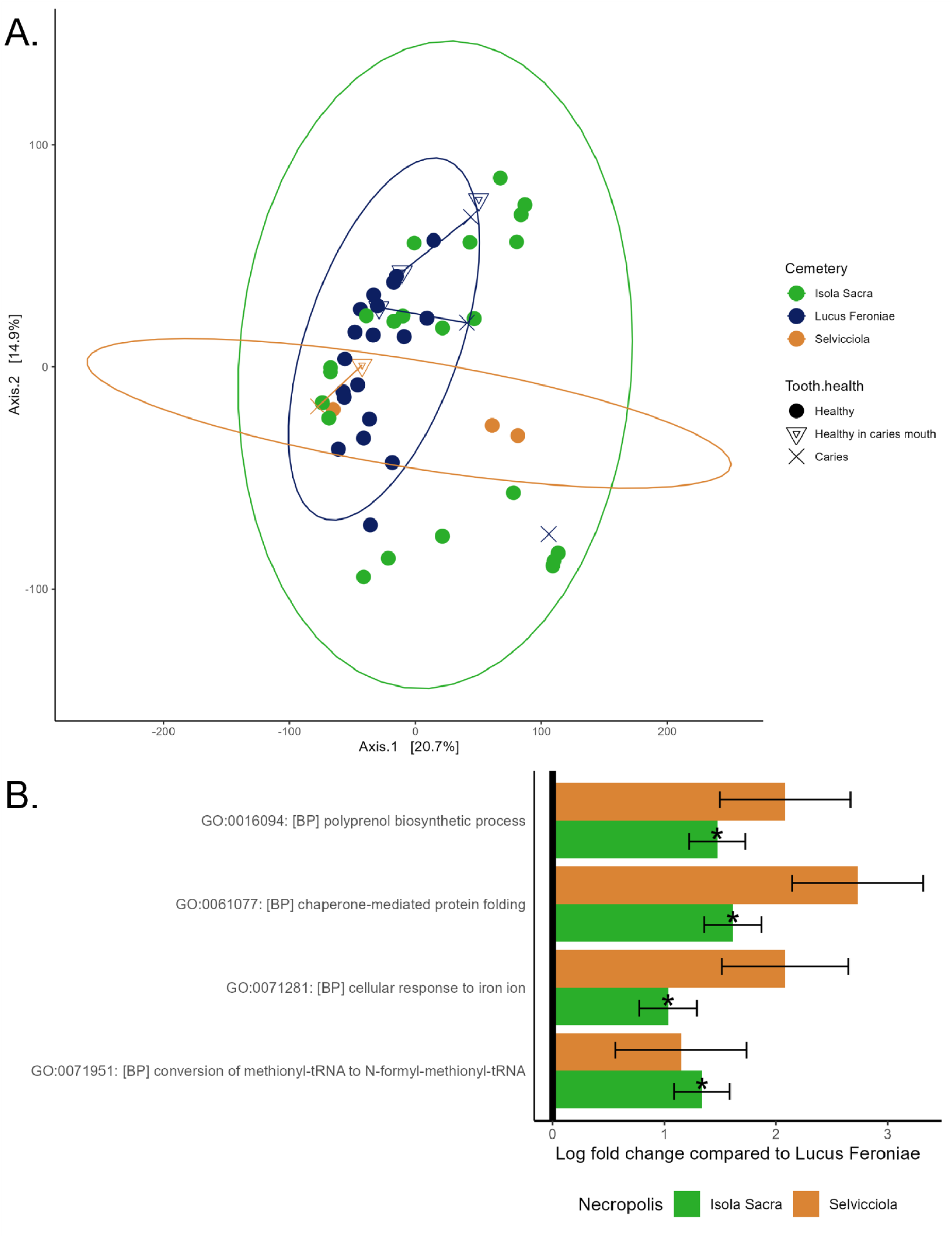
**A.** Functional profiles of the 52 retained dental calculus samples from Isola Sacra (in green), Lucus Feroniae (in blue) and Selvicciola (in orange) using CLR-transformed relative abundances of GO terms referring to biological processes. Three caries samples (cross symbols) and four samples of healthy teeth in individuals with caries (triangle symbols) are included. Three pairs of healthy-caries teeth from the same individual are connected by a line. The ellipses show the 95% confidence level using a multivariate t-distribution. **B.** Differentially abundant GO terms, as estimated by *ANCOMBC2*. The asterisks highlight processes that passed the sensitivity analysis for pseudo-count addition, always in the comparison with Isola Sacra. The error bars refer to the standard error of the log-fold change.

**Table 4.** PERMANOVA results for functional differences across the cemeteries, for 48 dental calculus samples after excluding poorly preserved and caries samples, showing the effect size (R^2^) and *p* value for read count (as a control for sequencing depth), health state (presence of caries in the sampled individual), and cemetery provenance for the dental calculus samples from Isola Sacra, Lucus Feroniae and Selvicciola based on Aitchison distance.

| **PERMANOVA terms** | **SumOfSqs** | **$R^2$** | **$p$ value** |
| --- | --- | --- | --- |
| Read count | 14489.000 | 0.067 | 0.003 |
| Cemetery | 17828.000 | 0.082 | 0.006 |
| Individual health state | 3665.000 | 0.017 | 0.526 |
| Residuals | 179721.000 | 0.830 |  |
| Total | 216620.000 | 1.000 |  |
| **Pairwise PERMANOVA** | **SumOfSqs** | **$R^2$** | **Adjusted $p$ value (Benjamini-Hochberg)** |
| Isola Sacra vs Lucus Feroniae | 13598.548 | 0.070 | 0.003 |
| Isola Sacra vs Selvicciola | 4714.071 | 0.030 | 0.601 |
| Lucus Feroniae vs Selvicciola | 6915.670 | 0.080 | 0.021 |

Four GO terms associated with biological processes were significantly enriched in Isola Sacra compared to Lucus Feroniae (GO:0016094: polyprenol biosynthetic process, GO:0061077: chaperone-mediated protein folding, GO:0071281: cellular response to iron ion and GO:0071951:conversion of methionyl-tRNA to N-formyl-methionyl-tRNA) (Figure 4B). These also appeared enriched in Selvicciola, however the comparison was not significant, possibly due to the small number of samples retained from this cemetery (N=4) (Figure 4B).

### Dietary analysis

After removing eukaryotic taxa with higher relative abundance in soil and control samples than in dental calculus samples (Table S6), we detected 30 plant genera, 23 mammalian genera and 17 fish species in the decontaminated and filtered dataset (Table S7). The implemented decontamination procedure allowed us to remove several contaminants and spurious hits, including plant species introduced into Europe from the Americas long after the study period and, hence, representing obvious false positives. However, despite these steps, one New World genus (*Sequoiadendron*) remained in the decontaminated plant dataset, and ten New World genera were present in the mammalian dataset, suggesting the retention of contaminant or spurious taxa (Table S6).

Each retained plant taxon showed, on average, 163.18 reads per sample (171.75 in dental calculus, 100.26 in soil and 1.39 in blanks, Table S8). All 30 plant taxa were detected in all three cemeteries. Five of them are plausible dietary taxa (*Brassica*, *Cornus*, *Prunus*, *Triticum* and *Vitis*), which were likely consumed by the individuals under study and still represent important food sources today (except *Cornus*). However, as these taxa were also detected in the co-analysed soil samples, suggesting their presence in the environment, direct consumption remains uncertain. The other detected genera were most likely not of dietary relevance but rather occurred in the environments surrounding the sites and cemeteries under study (e.g. *Juncus*, *Rhynchospora*), as they still occur in Italy today. Some taxa may have been used for timber (e.g. *Ilex*, *Populus*, *Quercus*) or for their medicinal properties (e.g. *Cardamine*, *Chenopodium*, *Trifolium*).

Each retained mammalian taxon presented, on average, 176.34 reads per sample (188.05 in dental calculus, 85.88 in soil and 1.08 in blanks). However, only 13 genera were likely present during Roman times, as the others are native to the Americas. Only two mammalian genera may have had dietary relevance in Italy (*Equus* and *Lepus*). As for the plant taxa, the mammalian genera detected could be naturally occurring in the environments around the archaeological sites, as they are also present in the soil samples.

Each retained fish taxon showed an average of 379.75 mapped sites per sample (427.18 in dental calculus, 704.92 in soil and 3.75 in blanks). All the species recovered, or their respective genera, may have been consumed as food, with the only exception of *Gymnothorax javanicus*, due to its potential toxicity (Dao et al., 2025), and *Doryteuthis pealeii*, likely not present in the region, as its native range is in the northwestern to central Atlantic. The species *Dicentrarchus labrax*, *Merluccius merluccius*, *Mullus surmuletus*, *Mytilus galloprovincialis* and *Sparus aurata* are present in the Mediterranean Sea and have been reported in the literature as having been consumed during Roman times (Marzano, 2013).

### Sex assignment

Of the 67 study individuals, 14 had ≥ 1,000 reads mapping to the X chromosome and the autosomes, and we therefore attempted molecular sex identification. Eleven samples were from dental calculus, whereas the remaining three samples were from skeleton-associated soils. These samples were collected by dislodging soil particles attached to human bones (Table S1) and are, therefore, likely to contain human DNA that leaked into the sediments during the decomposition period (Emmons et al., 2017). None of the soil samples had a paired dental calculus sample with a sufficient number of reads for molecular sex assignment.

We were able to assign molecular sex to seven individuals based on dental calculus samples and to all three soil-based samples (Table 5; Table S9). Sex assignment based on morphological features was available for eight individuals (Table 5) (Argenti & Manzi, 1988; Micarelli, 2020; Tafuri et al., 2018), allowing comparisons between molecular and morphological sex for six individuals. In four cases, morphological and molecular data were in agreement, whereas in two cases (one from dental calculus and one from soil) they disagreed (Table 5) (Argenti & Manzi, 1988; Tafuri et al., 2018). However, we note that the molecular sex assignment to individual RDC_2_H10 was less robust, with the likelihood ratio of X versus autosomes close to 0.8 (Table S9). The discrepancies between genetic and skeletal sex can be due to various factors, including the state of conservation of the material, contamination and insufficient number of human reads.

**Table 5.**
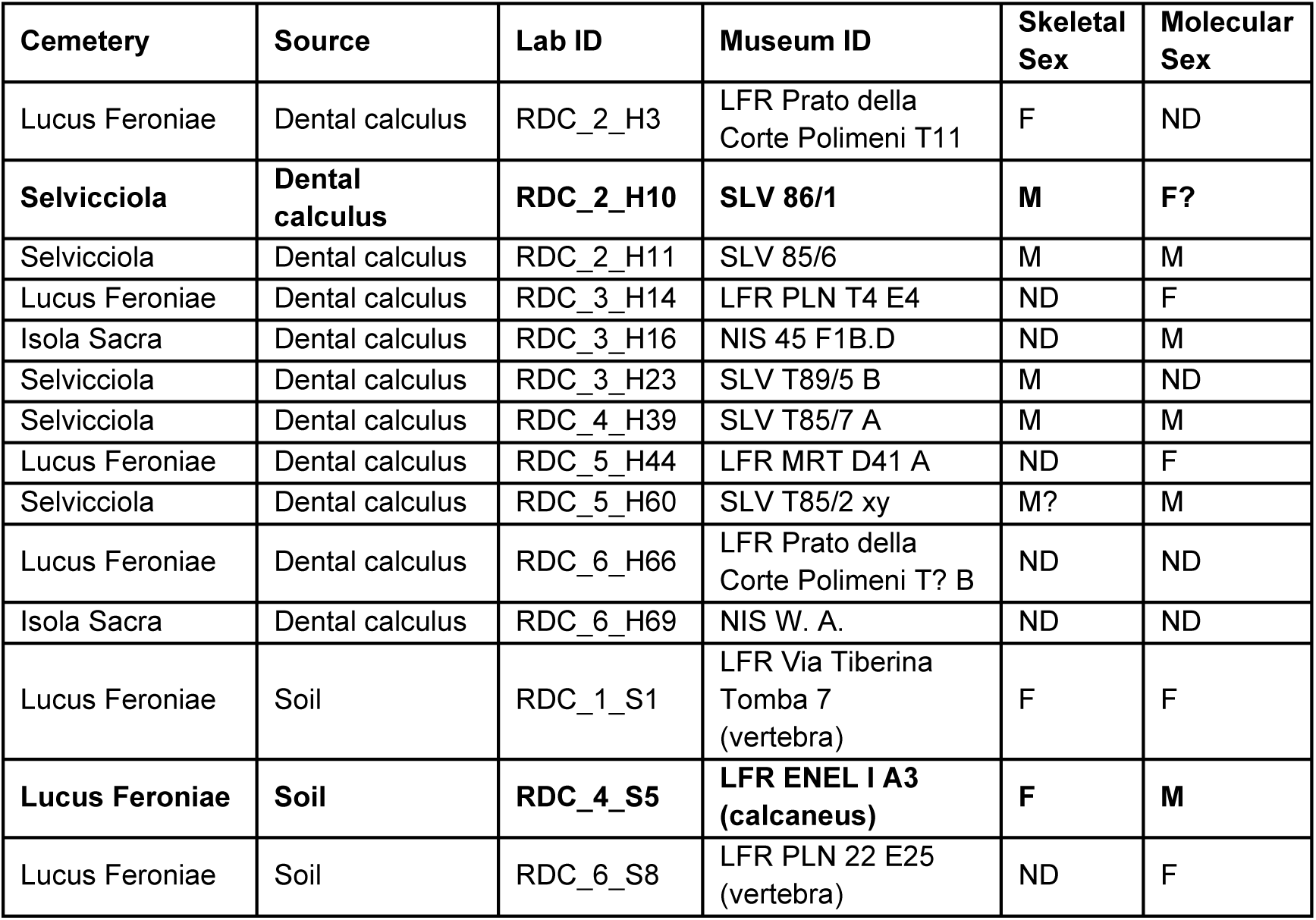
Skeletal and genetic sex assignments for 11 dental calculus samples and 3 soil samples. Bold entries depict discrepancies in sex identification between the methods. F = female, M = male, M? = probable male, F? = probably female, ND = not-determined.

### Geographic origins of the study individuals

Human dental calculus samples typically contain <1% eukaryotic reads, with the host read proportion being even lower (0.007% - 0.47%) (Mann et al., 2018). Nevertheless, we attempted to infer genetic affinities for samples for which at least 10,000 SNPs of the reference panel were covered by at least one sequencing read. Eleven samples satisfied this requirement and consistently showed the closest relationship to Tuscany/Iberia (Table S10). Other samples had too few sites for reliable population ancestry assignments. We are, therefore, unable to confidently infer any movement of individuals buried at the study cemeteries from the human genomic data contained in dental calculus.

## DISCUSSION

Our knowledge of the past is expanding through the combination of bioarchaeological methods applied directly to skeletal remains and the study of historical written records (Martin et al., 2013a, 2013b, 2013c), providing a more holistic view of ancient societies. Here, we used ancient DNA analysis of the microbiome preserved in dental calculus samples to enhance our understanding of the lifestyle of underrepresented social classes from the Classical and post-Classical periods. As farmers, freemen, slaves and war veterans are often overlooked in written records (Antonio et al., 2019; Cocozza et al., 2023; Cucina et al., 2006; Redfern, 2018; Wright et al., 2024), bioarchaeological methods provide a powerful (and often the only) tool to delve into their lives.

Over the last few years, dental calculus has gained recognition as an important and diverse source of information about past human populations (Bravo-Lopez et al., 2020; Eisenhofer et al., 2020; Gancz et al., 2023; Ottoni et al., 2021; Quagliariello et al., 2022; Velsko et al., 2022, 2023, 2024; Warinner et al., 2014). As it provides direct bioarchaeological evidence, the study of the oral microbiome can be applied to investigate the life of members of the society that might be “invisible”, rarely mentioned in historical records. In this study we analysed dental calculus samples from 67 individuals buried in three cemeteries around the city of Rome: Isola Sacra (I-III century CE), Lucus Feroniae (I-III century CE) and Selvicciola (IV-VIII century CE). This dataset allowed us to explore the oral microbiome, dietary choices and genetic provenance of these individuals, and to observe the transition from the peak of the Roman Empire to Late Antiquity. Our analyses reveal significant differences in the oral microbiome composition and function before and after the collapse of the Roman Empire, as well as between contemporary Imperial communities belonging to different social classes, suggesting the existence of marked differences among sections of Roman and post-Classical societies, possibly connected to temporal, geographic and socio-economic factors.

The oral microbial community composition and functional capacity of individuals buried in the cemetery of Lucus Feroniae differed significantly from both the contemporaneous Isola Sacra and the later-dated Selvicciola (Table 3, Table 4, Figure 3A, Figure 4A). Individuals buried in the rural town of Lucus Feroniae lived during a flourishing period of the Roman Empire, close to the city of Rome. Their rural inland location likely enabled them to grow food and provided reliable year-round access to it. In contrast, individuals buried in the urban coastal town of Isola Sacra, located on the direct commercial route towards the city of Rome, mostly obtained their dietary resources through trade and the income of food supplies might have been less stable. It is possible that this differential access to nutrition (De Angelis et al., 2020) is related to the differences in oral microbiome composition observed between the two cemeteries.

Our findings of significant differences between Lucus Feroniae and Isola Sacra agree with the results from previous studies. Although the archaeological record describes the inhabitants of Portus Romae as belonging to the middle class (Mannucci & Verduchi, 1996), palaeopathological analyses suggest sub-optimal health conditions and a relatively high incidence of physical stressors, visible on both teeth and long bones, indicative of poor eating habits during childhood and a more restricted access to resources, particularly when compared to individuals living in Lucus Feroniae (Manzi et al., 1989; Salvadei & Manzi, 1997). Portus was an important city located in a favourable area for commerce and was, therefore, densely populated. This might have created favourable conditions for disease to spread and might have led to frequent food crises, as experienced by several Roman cities (Garnsey, 1988). It is remarkable to observe differences in the taxonomic and functional composition of the microbiome between these communities that lived at the same time and were in close geographic proximity to each other but differed in social class. However, in non-human primates, subtle differences in ecology and diet affect the oral microbiome composition despite close geographic proximity (Moraitou et al., 2022). It is therefore conceivable that human societies will be similarly affected.

Following the collapse of the Roman Empire, the inhabitants of the Post-Classical site of Selvicciola could no longer rely on the centralised distribution system (e.g. the *annona* and *frumentatio*) and a stable self-production system might not yet have been established (Rotili, 2010). Late Antiquity was characterised by general political and social instability (Ledger et al., 2021). This likely had a negative impact on access to food and in turn impacted the oral microbiome composition, setting it apart from Lucus Feroniae. In this context, it is noteworthy that we do not find significant differences between Selvicciola and Isola Sacra, although they are as temporally distant from each other as Selvicciola and Lucus Feroniae. This suggests microbiome stability in populations experiencing potentially similar conditions regarding access to food resources before and after the collapse of the Roman Empire. It is, however, worth noting that only 11 dental calculus samples were collected from Selvicciola, four of which were excluded from the taxonomic analysis and additional two from the functional analysis due to poor preservation (Figure 2). Hence our results should be considered with caution. Based on previous studies of human dental calculus, these oral microbial communities are usually found to be remarkably stable through time, showing no or only very little differences in composition across considerably larger timescales than studied here (Gancz et al., 2023; Ottoni et al., 2021; Velsko et al., 2023), which is in line with the lack of differentiation between Selvicciola and Isola Sacra, despite the temporal difference between them.

Our results are in line with previous bioarchaeological analyses. Individuals buried in the cemetery of Selvicciola were found to be characterised by a poor health status when compared to both Isola Sacra and Lucus Feroniae, with a strong incidence of dento-alveolar lesions (Manzi et al., 1999; Salvadei & Manzi, 1997), especially caries, abscesses and antemortem tooth loss, and a high incidence of cribra orbitalia and cribra cranii, indicative of iron deficiency (Salvadei et al., 2001). Stable isotope analysis revealed that individuals buried in Selvicciola consumed a more restricted diet than those at the Classical site of Lucus Feroniae, likely because of a lower variability in protein sources (Tafuri et al., 2018). The low protein and high cereal consumption may explain the high incidence of health problems, caries and abscesses in Selvicciola, and may be linked to the differences in microbial composition we found in dental calculus between Lucus Feroniae and Selvicciola. Bioarchaeological tools, therefore, highlight a deterioration in the quality of life during a period of political decline.

Despite variations in the oral microbial community composition, we detected only a few differentially abundant microbial species in the oral microbiome from different cemeteries (Figure 3B,C). One such is *S. condimenti*, which was more abundant in Isola Sacra (Figure 3B,C) than in Lucus Feroniae. This bacterium is usually found in fish and fermented foods such as soy sauce (Gabrielsen et al., 2017). Its presence could be connected to the consumption of fish, as well as *garum* and *liquamen*, a fermented fish sauce and its byproduct, respectively, commonly used in Ancient Rome and often mentioned in *De Re Coquinaria*, a collection of recipes dated to the I-IV century CE. The higher abundance of *S. condimenti* observed in Isola Sacra could, therefore, be connected to the widespread consumption of fish and fish condiments among the individuals buried there. Being close to the Tyrrhenian coast, people in Isola Sacra might have produced or imported *garum* and *liquamen* to preserve their fish catches, having it more readily available than in Lucus Feroniae.

In terms of functional differences, Lucus Feroniae stands out by showing lower abundance of GO terms that are associated with stress responses. Specifically, functions related to polyprenol biosynthetic process (GO:0016094) and cellular response to iron ion (GO:0071281) are more abundance in Isola Sacra and Selvicciola (significantly so in the former) than in Lucus Feroniae (Figure 4B). Polyprenol is a lipid associated with peptidoglycan biosynthesis and cell responses to environmental stress (Surmacz & Swiezewska, 2011). Differences in genes involved in iron ion biosynthesis may reflect the degree of exposure to this element, either through diet or from the environment.

Although the differences we observe between the cemeteries may result from differences in diet and nutrition, we are unable to provide evidence for this. We identified the same plant, mammal and fish taxa in all three cemeteries. As previously demonstrated, extracting dietary information from dental calculus is riddled with challenges (Mann et al., 2023). Plant and animal DNA is considerably less abundant than microbial DNA, eukaryotic databases are not as comprehensive as those used for bacterial and archaea classification, and eukaryotic genomes are usually considerably larger than prokaryotic genomes, increasing the rate of spurious mappings and thereby reducing the information content from the few mapped reads (Mann et al., 2023; Oskolkov et al., 2025). In addition, there is a general difficulty to distinguish dietary taxa from environmental contamination. While some of the taxa detected in dental calculus samples from this study could plausibly represent dietary items (*Equus*, *Lepus*, *Brassica*, *Cornus*, *Prunus*, *Triticum* and *Vitis*), others are likely to be naturally occurring around the sites and the cemeteries both during and after the lifetime of the studied individuals. All eukaryotic taxa that passed our filters were also detected in soil samples. Therefore, even after decontamination, we cannot exclude that the individuals came into contact with the eukaryotic species only after death. At the same time, it is not surprising to detect signals of cultivated plants and farmed animals in soil samples, given that they might have been planted and cared for in the same place humans lived and consumed them. In summary, disentangling dietary and environmental signals from dental calculus remains a considerable challenge, which can likely only be solved by consulting additional sources of information on food consumption in the past (such as written records, organic residues in ceramic vessels and artistic depictions).

Because dental calculus also contains some host reads, we attempted to infer the provenance of the individuals under study. This is of particular interest in the context of the Roman Empire, which has been associated with high population mobility. Only a few studies are available for Italy during the Roman period, which depict a cosmopolitan society characterised by various genetic ancestries (Antonio et al., 2019; Ravasini et al., 2024). In Lucus Feroniae, historical sources point to a diverse population, including war veterans, slaves and freemen from various regions of the Empire. However, only eleven individuals yielded sufficient data to attempt ancestry inferences, and all of them were consistent with close genetic similarity to present-day Tuscany/Iberia. Overall, without applying a targeted enrichment protocol (Ozga et al., 2016), the number of endogenous reads was too low for meaningful population-level analyses. These findings are in line with previous studies highlighting the limited presence of host DNA in human dental calculus samples (Mann et al., 2018), in contrast to the considerably higher proportion of host DNA in dental calculus from non-human mammals (Brealey et al., 2020; Moraitou et al., 2022, 2025).

Despite very low sample sizes, our analyses suggest differences in microbiome composition, but not alpha diversity, between samples from healthy and carious teeth (Figure 3A, Supplementary Figure S3). The composition of microbial communities from carious teeth does not only seem to be different from healthy teeth within the same individual but also tends to occupy peripheral positions in the PCoA space of all considered samples (Figure 3A). While dedicated studies with larger sample sizes are needed to understand the composition of these diseased oral communities in ancient human societies and to compare them with present-day pathologies, the observed differences are in line with the Anna Karenina principle for dysbiotic microbiomes (Zaneveld et al., 2017).

## CONCLUSIONS

By studying the oral microbiome of individuals buried in three different cemeteries spanning from the I to the VIII centuries CE, we detected differences between both contemporary and asynchronous populations, which could result from differences in subsistence practices, access to resources and general health status. To the best of our knowledge, we present the first record of differences in oral microbiome taxonomic and functional composition in human societies that is associated with social class, possibly as a result of trade, local farming practices and population density. Further analyses are needed to better understand dietary choices at the three sites and how they might have impacted oral microbial communities. Overall, we find a good correspondence between the results obtained from dental calculus and previous palaeopathological, morphological and isotopic studies, thereby providing further resolution to our knowledge of past human societies, particularly for previously underexplored groups.

## ACKNOWLEDGEMENTS

This study has been funded by the Italian Ministry of University and Research (MUR) under the Programma Operativo Nazionale Ricerca e Innovazione (PON R&I) 2014–2020 (FSE REACT-EU) programme (Grant agreement No. DOT1326JZS to M.F.) and by the Swedish Phytogeographical Society (“Lundman Foundation for Anthropological Studies” scholarship, years 2022 and 2023, to M.F.). We would like to thank Elisabetta Asloisi Masella (technical assistant of the Museum of Anthropology “Giuseppe Sergi”, Sapienza University of Rome) for her help during sample collection. We would also like to thank Dr. Paola Francesca Rossi (official at the Archaeological Park of Ostia Antica) for her useful comments on the manuscript draft. We also thank Gunilla Engström (Animal Ecology Department, Uppsala University) for her help in the laboratory. Sequencing was performed by the SNP&SEQ Technology Platform in Uppsala, which is part of the National Genomics Infrastructure (NGI) Sweden and Science for Life Laboratory. The SNP&SEQ Platform is also supported by the Swedish Research Council and the Knut and Alice Wallenberg Foundation. The computations were enabled by resources in projects UPPMAX 2025/2-463, NAISS 2025/5-646, and NAISS 2025/6-415 provided by the National Academic Infrastructure for Supercomputing in Sweden (NAISS) at UPPMAX, funded by the Swedish Research Council through grant agreement no. 2022-06725.

## DATA AVAILABILITY STATEMENT

The data that support the findings of this study are openly available in the European Nucleotide Archive (ENA) at https://www.ebi.ac.uk/ena/browser/home, reference number PRJEB106785.

