## Supplementary Figures 1-4 for "Lifestyle impacts the oral microbiome of Classical and Post-classical societies in Italy"

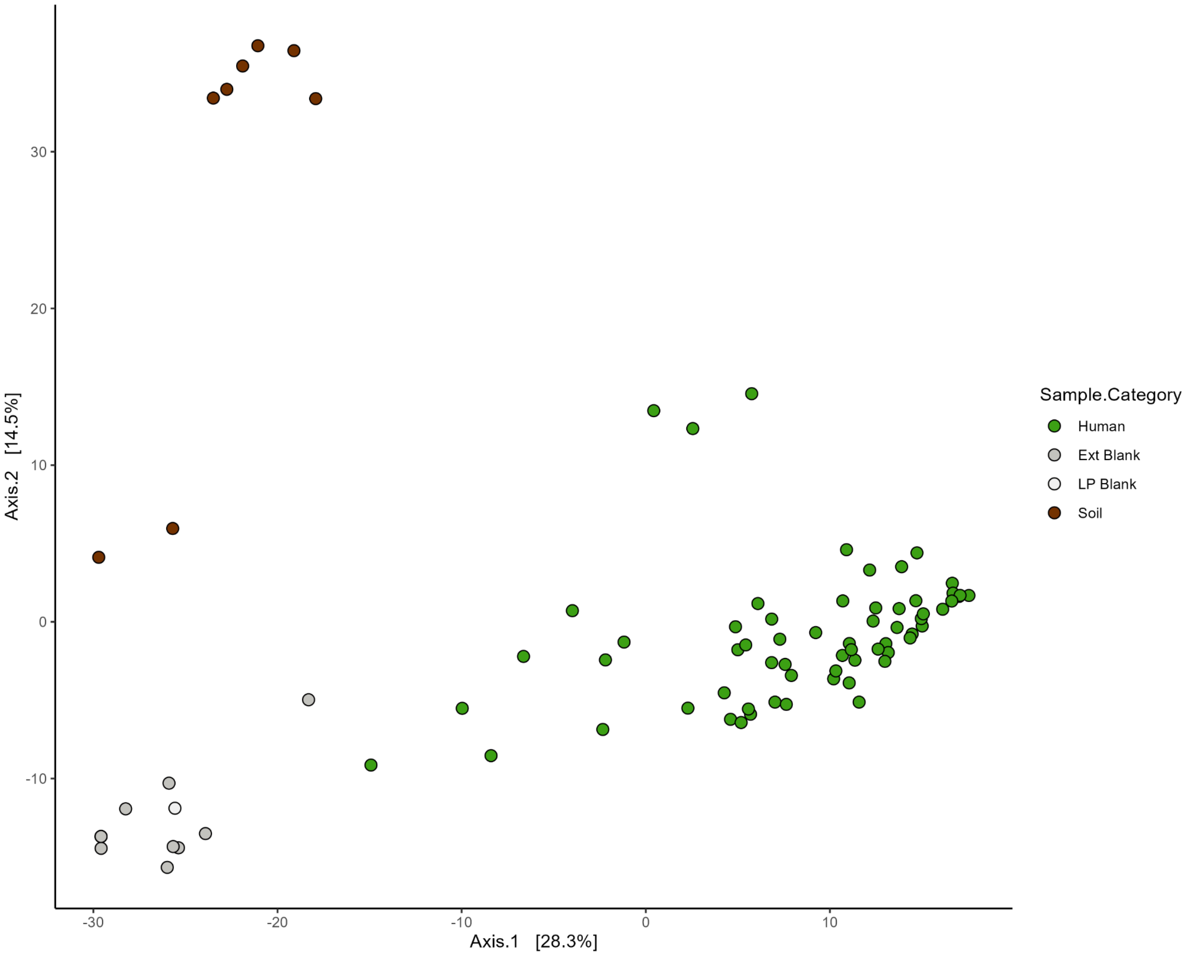


**Supplementary Figure S1.** Principal Coordinate Analysis (PCoA) ordination based on the taxonomic composition of dental calculus (in green), extraction and library blanks (in dark and light grey, respectively) and soil (in brown) following decontamination.

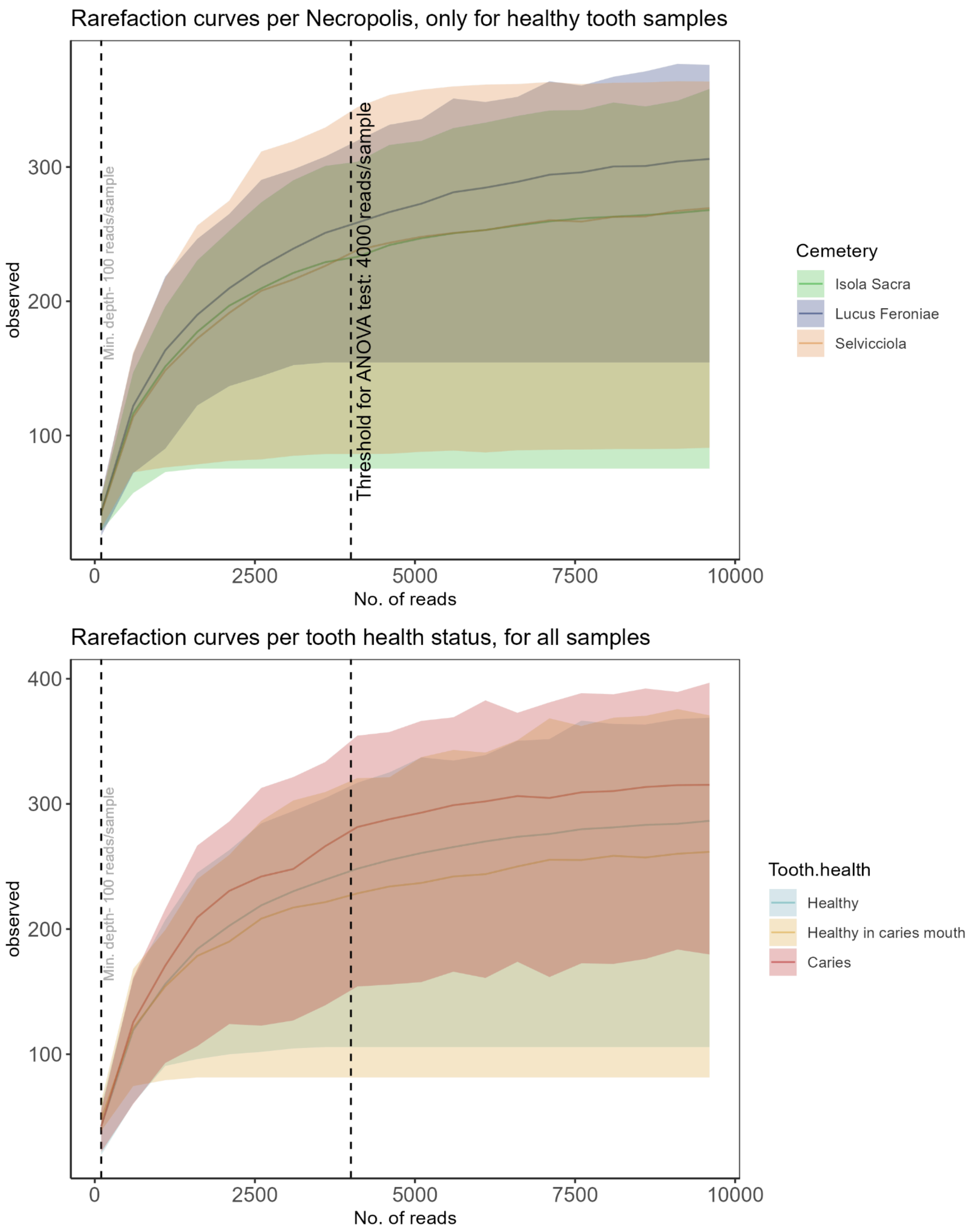


**Supplementary Figure S2.** Rarefaction curves showing observed microbial diversity when subsampling each sample to 100-10,000 reads. **A.** Only healthy teeth, coloured by cemetery of origin. **B.** All samples, coloured by health state.

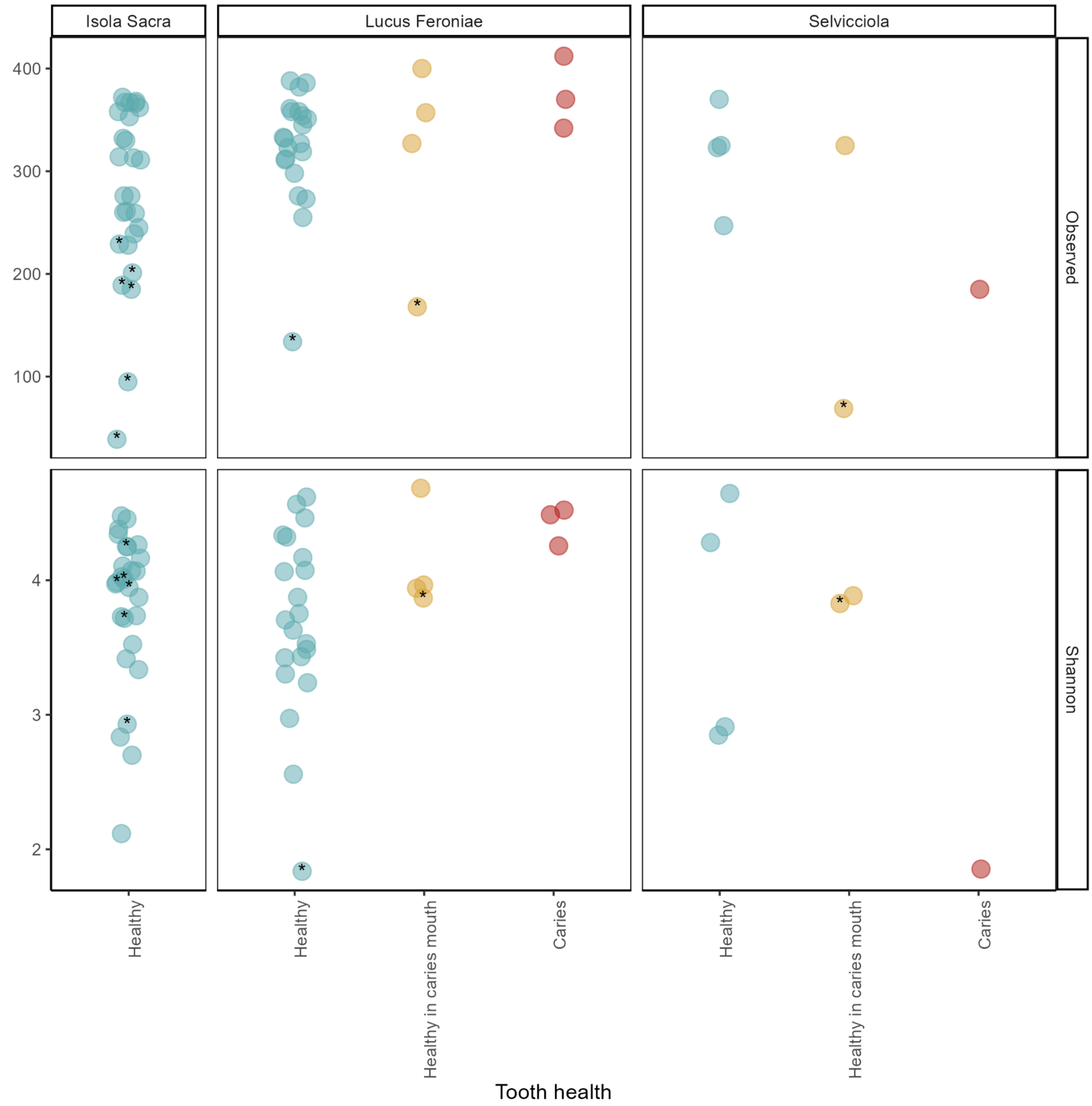


**Supplementary Figure S3**. Observed species richness (top row) and Shannon evenness index (bottom row) for different cemeteries and tooth health states. Asterisks indicate samples that were excluded from alpha diversity tests with ANOVA because they had fewer than 4,000 taxonomically classified reads after filtering and decontamination. “Healthy in caries mouth” corresponds to dental calculus microbiome from healthy teeth sampled in an oral cavity with presence of caries elsewhere, whereas “Caries” samples were obtained directly from the caries lesions.

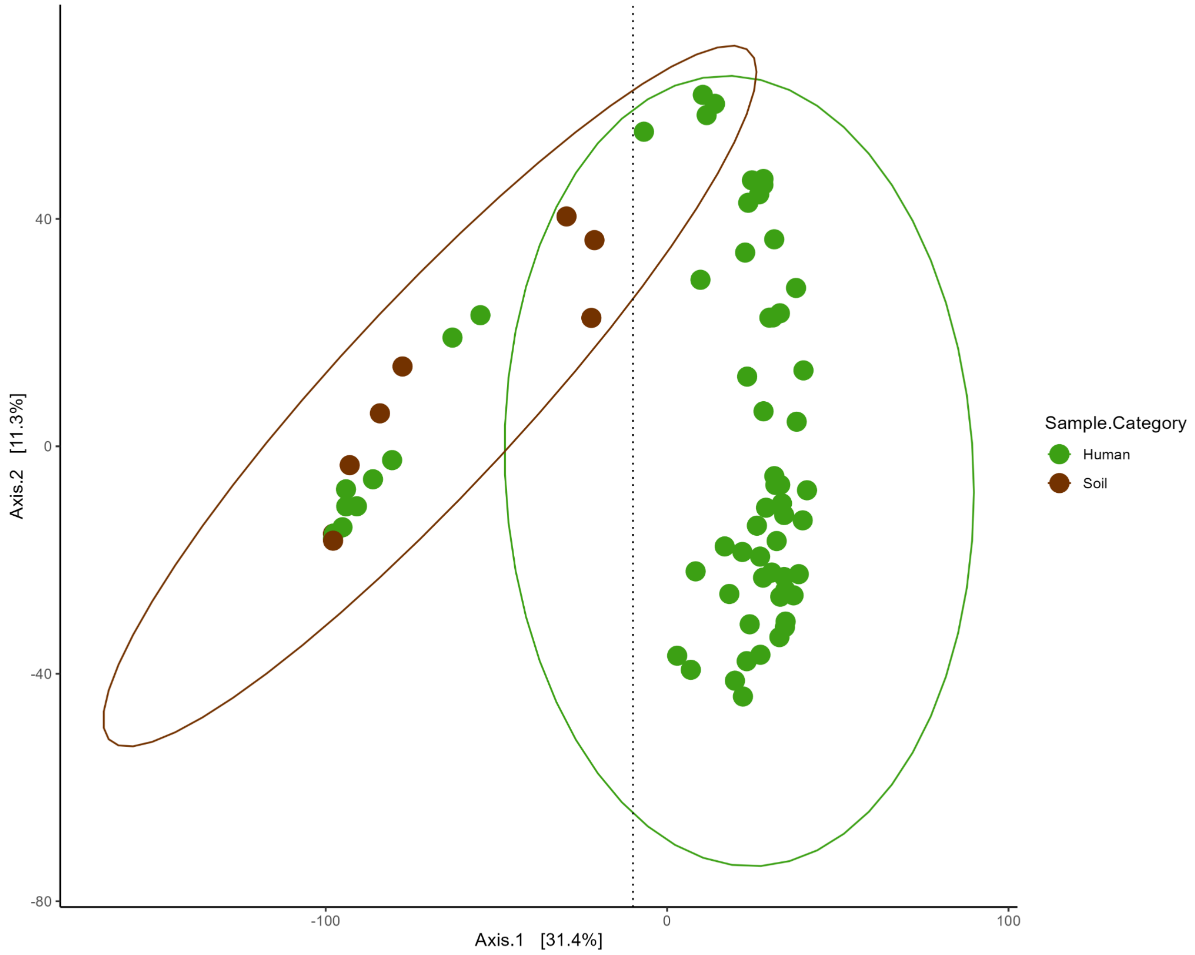


**Supplementary Figure S4**. Principal Coordinate Analysis (PCoA) ordination based on the GO term composition (only biological processes) of dental calculus (in green) and soil (in brown). The ten dental calculus samples on the left of the dotted line were considered poorly preserved, as they clustered closer to the soil samples, and were consequently excluded from downstream functional analysis.
